# Enhanced leaky sex expression is an adaptive plastic response to pollen limitation in the dioecious plant *Mercurialis annua*

**DOI:** 10.1101/703009

**Authors:** Guillaume G. Cossard, John R. Pannell

## Abstract

Phenotypic plasticity is almost ubiquitous across the tree of life, but clear demonstrations that it is adaptive are rare. In dioecious plants, males and females frequently show ‘leaky’ sex expression, with individuals occasionally producing flowers of the opposite sex. Here, we demonstrate that leaky sex expression in the wind-pollinated dioecious herb *Mercurialis annua* is plastically responsive to its mating context. We compared experimental populations of females growing either with or without males. Females growing in the absence of males were leakier in their sex expression than controls with males, producing more than twice as many male flowers. Moreover, because greater leakiness was more highly represented in the population’s progeny, we conclude that enhanced leakiness in sex expression is adaptive in *M. annua*. We discuss differences in the degree of plasticity between ploidal races of *Mercurialis annua* in terms of likely differences in the reliability of the signal plants may perceive for the presence of males in their populations. Our results provide a striking instance of adaptive plasticity in the reproductive behavior of plants and draw attention to possible constraints on plasticity when the environmental signals that individuals receive are unreliable.

## Introduction

The expression of an individual’s phenotype will commonly depend on its environmental context (Schlichting and Pigliucci 1998; Pigliucci 2001). Such phenotypic plasticity can have several causes (Van Kleunen and Fischer 2005; Josephs 2018). First, it may be a passive response to different growth conditions, such as variation in resource availability that limits the size of individuals or their capacity to invest in particular structures (Sultan 1987; Dorn et al. 2000; Van Kleunen and Fischer 2005). Second, phenotypic plasticity may be the result of developmental programs that are altered by exposure to unaccustomed stress in novel environments, such as excessive heat, toxicity or physical damage (Van Kleunen and Fischer 2005). Third, phenotypic plasticity may be an adaptive expression of genotypes habitually exposed to heterogeneous environments (Gomulkiewicz and Kirkpatrick 1992), such that plastic genotypes have greater fitness, averaged over the environments they may encounter, than non-plastic ‘canalized’ genotypes that always express the same phenotype (Releya 2002; Van Kleunen and Fischer 2005).

Although phenotypic plasticity is all but ubiquitous, plants are particularly plastic organisms (Bradshaw 1965; Sultan 1987; Josephs 2018). This is likely due both to their sessile nature, which means that they cannot actively choose their habitat and must make the best of the conditions to which they are exposed, as well as to their modularity, which allows individuals to modify the phenotype of their modules (e.g., branches, inflorescences or flowers) as they are produced in response to environmental variation over time and space. Nevertheless, evidence that plastic variation in plants is adaptive remains surprisingly thin, with few clear demonstrations that the different phenotypes expressed in different environments actually improve fitness (Van Kleunen and Fischer 2005; Hendry 2015; Wagner and Mitchell-Olds 2018). Several studies have demonstrated that plant responses to shading are adaptive, with fitness benefits to individuals that produce longer internodes and achieve greater height when overtopped by or growing beside potential competitors (e.g., Dudley and Schmitt 1995, 1996). The plastic induction of increased defense in response to herbivory has also been shown to increase individuals’ fitness (e.g., Karban et al. 1997; Agrawal 1998; Agrawal 1999). Further, Baythavong (2011) and Baythavong and Stanton, (2010) showed that variation in a number of morphological and phenological traits was adaptive in environments with small-scale variation in soil chemistry, and Kenney et al. (2014) showed that plasticity in water-use efficiency (WUE) conferred fitness benefits under dry conditions – though Nicotra and Davidson (2010) reported inconsistent findings in their review of plasticity in WUE among studies.

Plant reproductive traits may also be phenotypically plastic. Examples include: variation in reproductive effort or reproductive allocation, which is sensitive to resource availability and competition (Weiner 2004); sex allocation, e.g., in terms of the relative numbers of male versus female flowers produced by monoecious individuals, which varies with plant size and resource status (Pannell 1997; Dorken and Barrett 2003; Paquin and Aarssen 2004); and floral longevity and floral display size, with plants adjusting their attractiveness to pollinators in response to the relatedness of their neighbors (Torices et al. 2018) or as a function of relative pollinator abundance and visitation rates (Harder and Johnson 2005). This latter example is particularly interesting in the context of our study here, because it indicates the extent to which plants may alter their reproductive allocation decisions specifically in response to plant mating opportunities. Specifically, Harder and Johnson (2005) found that floral display in the hermaphroditic orchid *Satyrium longicauda* was enhanced when pollinator visitation rates were low, increasing the possibility for later pollen receipt or geitonogamous self-pollination, with likely fitness benefits. Similarly, Lopez et al. (2003) found that in the monoecious plant *Begonia gracilis*, individuals whose female flowers were pollen-limited produced more male flowers than those whose female flowers enjoyed experimentally augmented pollen deposition, suggesting that plants can respond to the operational sex ratio of the population (though without demonstrating a clear effect on fitness in natural populations). In homosporous ferns, gametophytes are more likely to develop as males when females or hermaphrodites are locally abundant, a switch mediated by interplant chemical signaling (Banks 1997) and interpreted as an adaptive response to mating opportunities (Banks 1997).

Many angiosperms with separate sexes also show variation in sex expression. Specifically, the males and females of dioecious plants commonly display inconstant or ‘leaky’ sex expression, with the occasional production of a few flowers of the opposite sex (e.g., Baker 1967; Lloyd 1972; Lloyd and Bawa 1984; Diggle 1991; Korpelainen 1998; Delph 2003; Venkatasamy et al. 2007). Such leaky sex expression, which is more common in males than females (Delph and Wolf 2005; Ehlers and Bataillon 2007; but see Cossard and Pannell 2019), has probably been important in facilitating evolutionary transitions from dioecy to hermaphroditism under conditions of mate limitation (Ehlers and Bataillon 2007; Crossman and Charlesworth 2014; Käfer et al. 2017). It is plausible that leaky dioecy may be adaptive by assuring reproductive success under pollen- or mate-limited conditions (Sultan 1987; Korpelainen 1998; Ghalambor et al. 2007), e.g., in colonizing species, whose population densities (and consequently whose mate availability) may vary greatly; colonizing species are thought to be particularly likely to benefit from plastic strategies in general (Baker 1965; Stebbins 1965; Sultan and Spencer 2002). However, to our knowledge the adaptiveness of leaky dioecy has not been demonstrated for any dioecious plant population. Indeed, leaky sex expression can be elicited by external stimuli such as temperature, drought, simulated herbivory or exogenous hormone application (Kuhn 1939; Westergaard 1958; Korpelainen 1998; Delph and Wolf 2005; Golenberg and West 2013), none of which suggest an obvious adaptive function. Importantly, there appears to be no empirical support to date for the possibility that leaky sex expression might be prompted by pollen or mate limitation, which would be more plausibly adaptive.

Here, we demonstrate that leaky sex expression in the dioecious, wind-pollinated annua herb *Mercurialis annua* is plastic, and that the expression of enhanced leakiness under conditions of altered mate availability is adaptive. Dioecious *M. annua* has an XY system of sex determination in which the Y chromosome has a mildly degenerate non-recombining region (Li et al.; Veltsos et al. 2018; Veltsos et al. 2019). Sex ratios in wild populations are typically 1:1 (Russell and Pannell 2015). The species is strongly sexually dimorphic, with males and females differing in a number of physiological, life-history and morphological characters (Sanchez-Vilas and Pannell 2010; Hesse and Pannell 2011b; Sanchez-Vilas and Pannell 2011; Sanchez-Vilas et al. 2011; Labouche and Pannell 2016; Tonnabel et al. 2017). Yampolsky (1919; 1922) noted the presence of “intergrades” in both sexes of *M. annua* (evidently individuals showing leaky sex expression), though Cossard and Pannell (2019) showed that females are more often leaky than males. Yampolsky (1930) and Kuhn (1939) demonstrated that leakiness in *M. annua* could be enhanced by pruning, but it was not obvious from these studies that the plastic response was adaptive rather than simply a physiological response to unaccustomed stress (Van Kleunen and Fischer 2005).

Our experiment involved growing females of *M. annua* in populations with or without males. We hypothesized that, under a scenario of adaptive leaky sex expression, females growing without males would be more likely to produce male flowers, and would produce more of them. We then verified that leakiness in the absence of males is adaptive in *M. annua* by demonstrating an evolutionary response to selection on enhanced leakiness in the progeny generation. There are several reasons to expect that selection might have favored a plastic leakiness in sex expression in response to variation in mate availability. First, the dioecious populations of *M. annua* are known to have expanded their range recently from the eastern Mediterranean Basin into western Europe (Obbard et al. 2006; Gonzalez-Martinez et al. 2017), during which populations establishing at the colonizing front are likely to have been exposed to mate-limited conditions. Second, the species has a metapopulation structure and dynamic, with frequent population turnover and substantial fluctuations in population size from generation to generation (Eppley and Pannell 2007a; Dorken et al. 2017), conditions under which a leaky expression of the opposite sex would likely confer fitness benefits on an on-going basis, even after a range expansion ended (Golenberg and West 2013). Third, previous work has shown that females of *M. annua* quickly become pollen-limited at low density (Hesse and Pannell 2011a). Fourth, inbreeding depression in western European populations is low (Eppley and Pannell 2009), perhaps as a result of the range expansion (Pujol et al. 2009; Gonzalez-Martinez et al. 2017), so that selfing by leaky individuals under pollen limitation would seem to be particularly likely to be beneficial (Wolf and Takebayashi 2004; Pannell 2008). Finally, individuals in female-only populations established by leaky females would have particularly high siring success if they could respond to the absence of males by producing more pollen (see Dorken and Pannell 2008, 2009).

## Material and methods

We established six experimental populations of dioecious *Mercurialis annua* in separate common gardens on the campus of the University of Lausanne and in gardens around the city. Three ‘control’ populations were established at a 1:1 sex ratio (90 males and 90 females), and three ‘female-only’ population comprised only females (180 females). Plants were established for the experiment from seeds pooled from 35 demes of a metapopulation in north-western Spain (Labouche and Pannell 2016), and seedlings were first raised for six weeks in a glasshouse before being transplanted into their experimental plots. After seven weeks of subsequent growth, we recorded the number and mass of male flowers and seeds for 35 to 50 females per population, and calculated the male and female reproductive efforts (MRE and FRE) by dividing the total male-flower mass and seed mass, respectively, by the total above-ground biomass.

We analyzed the expression of leaky sex expression by females in terms of (1) the proportion of leaky females in the population and (2) their average MRE. To calculate proportions, we defined a leaky female as one with an MRE greater than the 95 percentile MRE across the control populations. By this definition, an average 5% of females were identified as leaky across the three control populations. We chose the 95% cut-off for our definition because it can be easy to miss one or two small male flowers on a large plant, and for coherence with other work on leaky sex expression in the lab (Cossard and Pannell 2019). An analysis based on an absolute measure of leakiness (including all females with any male-flower production at all) yielded qualitatively similar results.

We compared the proportion of inconstant individuals between female-only and control populations using a generalized linear model with a binomial error distribution, and with population replicates included as a random factor. MRE and FRE were compared using permutation *t*-tests, with 100,000 bootstraps and two degrees of freedom (i.e., with population replicate as the unit of observation).

To assess whether enhanced leaky sex expression conferred a fitness advantage on females growing in the absence of males, and thus to test whether it was adaptive, we sought evidence for an evolutionary response to selection in the progeny generation. While an observed evolutionary response to selection towards enhanced leakiness in the progeny of parents growing without males demonstrates a heritable component to leaky sex expression over and above any plastic component, it also provides unambiguous evidence that parents with enhanced leakiness were fitter under those conditions. We use a simple model to demonstrate that such a response satisfies the minimal conditions for concluding that the plasticity we see must be adaptive, even under the conservative scenario where reduced leakiness in the presence of males confers no advantage.

## Results

There was a tendency for leaky females to be more frequent in female-only than control populations, but the difference was not significant (Figure 1a; *P* = 0.16). Nor could we detect any significant difference in the FRE between the two treatments (*P* = 0.35), although variance of FRE was greater in male-deprived populations (*F*-test *P* = 7.06 ×10^−7^; variance = 1.19 ×10 ^−3^ and 4.83 ×10^−4^ for female-only and control populations, respectively; Figure 1c). In contrast, the MRE of females in female-only populations was 2.33 times that of females in control replicates (*P* = 4.71 ×10^−3^; Figure 1b).

**Figure 1.**
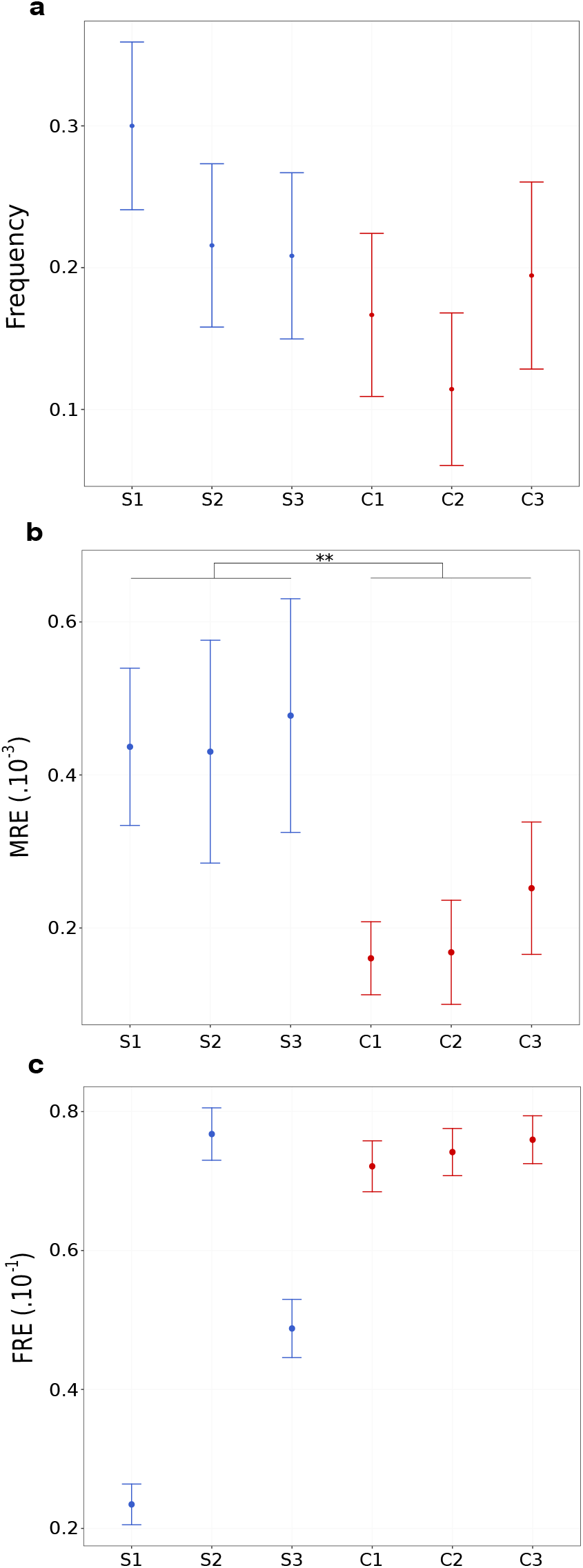
(a) Proportion of females with leaky sex expression, (b) male-reproductive effort (MRE) of females, and (c) female reproductive effort (FRE) of females in male-only (blue) and control populations (red). Means and standard errors are shown. *\*\**: 0.001 < *P* < 0.01.

## Discussion

Our results provide evidence for a plastic component to the expression of sex inconstancy in *M. annua*, with a slight tendency for more leaky females in populations lacking males and, significantly, females that on average produced more than twice as many male flowers in the absence of males than in their presence. Plastic responses to pollen limitation in plants have been recorded previously, in the context of the increased production of cleistogamous flowers and corresponding selfing rates in *Viola praemorsa* (Jones et al. 2013) and *Collomia grandiflora* (Albert et al. 2011). However, although leaky sex expression is a common feature of the reproduction of dioecious plants (Lloyd and Bawa 1984; Korpelainen 1998; Ehlers and Bataillon 2007; Cossard and Pannell 2019), our study demonstrates that such leakiness may be plastic and sensitive to mating opportunities. It is too early to speculate on the frequency of adaptive plasticity in leaky sex expression in plants more generally, but given that mate availability likely varies substantially in natural populations, it would seem to be a trait that dioecious populations should often evolve.

*M. annua* is a ruderal species that occupies disturbed habitat, and previous work has shown that its populations are subject to substantial fluctuations in population size and population turnover, with local extinctions and colonization by seed dispersal being key features of the species’ ecology (Pannell 1997; Eppley and Pannell 2007b; Dorken et al. 2017). In sparse populations of *M. annua*, female reproduction may be strongly pollen-limited (Hesse and Pannell 2011a), and a capacity to produce male flowers and to self-fertilize under these conditions is likely to be adaptive, especially as inbreeding depression is low (Eppley and Pannell 2009). It is thus plausible that plasticity in leaky sex expression has evolved and/or has been maintained under conditions of fluctuating population size and density in metapopulations. *M. annua* came to occupy its broad range in western Europe via a recent range expansion from the eastern Mediterranean Basin (Obbard et al. 2006; Gonzalez-Martinez et al. 2017), and it is plausible that the repeated demographic bottlenecks that occurred during the range expansion also favored the maintenance of plasticity in leaky sex expression.

Ultimately, plasticity in sex expression in dioecious *M. annua* should be maintained as long as its benefits outweigh its costs. Potential costs of plasticity, which might constrain its evolution (reviewed in Van Kleunen and Fischer 2005), include that of acquiring accurate information about the state of the environment, and that of maintaining the machinery required to sense and respond to environmental cues. Despite considerable effort, however, it has been difficult to find any evidence for such costs (Auld et al. 2010). Rather, it seems more likely that the evolution of plasticity is held in check by so-called ‘limits’ associated with its deployment, such as the penalty paid by individuals that respond incorrectly to an unreliable signal, or the disadvantage of having to wait until a signal is perceived before expression of an appropriate phenotype (Auld et al. 2010; Murren et al. 2015). We do not know which of these limits might apply to plasticity in leaky sex expression, but both seem plausible. In short-lived species like *M. annua*, or in species with a short reproductive season, there might be strong disadvantages associated with delaying leaky sex expression until mating prospects have been perceived. Yet such delays seem inevitable if, as seems likely, the signal to which plants are responding is the actual deposition of pollen on their stigmas, or physiological signals arising from seed-filling or fruit-set that follow pollination.

Regional variation in the plasticity of sex expression in *M. annua* hints at possible limits to its adaptive expression in terms of signal reliability. Although leaky sex expression is plastic in dioecious *M. annua*, as this study shows, polyploid populations of the species in the Iberian Peninsula comprise females that always express a male function, i.e., they are canalized in their sex expression as monoecious individuals. The reason for this difference is not immediately clear, because the two lineages are both colonizers of very similar disturbed habit (Buggs and Pannell 2007) and might both benefit from the plasticity we have reported here. However, whereas the absence of males from a population in the dioecious diploid lineage would be reliably perceived by females in terms of an absence of pollen on their stigmas, or low rates of fertilization of their ovules, monoecious ‘females’ of the polyploid lineage would receive pollen from other monoecious individuals even when males are absent, which occurs frequently (Pannell et al. 2014). Thus, although polyploid monoecious individuals would likely benefit from suppressing the production of male flowers when males are present, because males are so much better at dispersing pollen and outcross siring (Eppley and Pannell 2007a; Santos del Blanco et al. 2018), this strategy might be inaccessible to them for want of a suitably reliable signal.

In conclusion, our study has demonstrated that leaky sex expression in *M. annua* has a substantial plastic component, and our discussion above suggests that this strategy is likely adaptive. Consistent with this adaptive interpretation, Cossard et al. (2019) have recently shown that females of *M. annua* deprived of their mates, as in the female-only lines here, are indeed under strong positive selection for enhanced leaky sex expression, and respond to this selection by evolving enhanced leakiness. Our conjecture that plastic leakiness in sex expression in *M. annua* is adaptive would thus seem to be well-founded. Leaky sex expression in dioecious populations is almost certainly a necessary requirement for evolutionary transitions to occur from dioecy to hermaphroditism (Ehlers and Bataillon 2007; Crossman and Charlesworth 2014; Käfer et al. 2017). While we might expect such transitions to be slowed if leaky expression is strongly plastic, because plasticity diminished a trait’s heritability (Falconer and Mackay 1996), it is tempting to speculate that environmentally induced induced leakiness, as found in diploid *M. annua*, might actually enhance the potential for the evolution of canalized monoecy or hermaphroditism via genetic assimilation (Waddington 1942, 1953, 1959; Pigliucci and Murren 2003; Price et al. 2003; West-Eberhard 2003; Ghalambor et al. 2007; Schlichting 2008).

## Acknowledgements

We thank J. Gerchen and T. Martignier for comments on the manuscript. The research was funded by the Swiss National Science Foundation.

